# Functional plasticity and reversible growth behavior of patients’ acute myeloid leukemia stem cells growing in mice

**DOI:** 10.1101/707935

**Authors:** Sarah Ebinger, Christina Zeller, Michela Carlet, Daniela Senft, Wen-Hsin Liu, Maja Rothenberg-Thurley, Klaus H. Metzeler, Karsten Spiekermann, Binje Vick, Irmela Jeremias

## Abstract

Resistance against chemotherapy remains a major obstacle in treating patients with acute myeloid leukemia (AML). Novel therapeutic concepts are especially desired to target and eliminate resistant AML stem cells. Here we show that AML stem cells harbor the plasticity to switch from a low-cycling, chemotherapy resistant state into an actively proliferating state associated with treatment response. We used patient-derived xenograft (PDX) cells from patients with high risk or relapsed AML, which were lentivirally transduced for marker expression, stained with the proliferation-sensitive dye Carboxyfluorescein succinimidyl ester (CFSE), and re-transplanted into next-recipient mice. A rare subpopulation of AML cells displayed reduced proliferation *in vivo*, associated with increased treatment resistance. The proportion of AML cells with stem cell potential was identical in both, the highly and lowly proliferative sub-fraction. In re-transplantation experiments, proliferation behavior proved reversible, and AML stem cells were able to switch between a high and low proliferation state. Our data indicate that AML stem cells display functional plasticity *in vivo*, which might be exploited for therapeutic purposes, to prevent AML relapse and ultimately improve the prognosis of patients with AML.

## Introduction

AML patients are at risk to suffer disease relapse associated with dismal prognosis (1). The rare subpopulation of AML stem cells (or LIC for leukemia initiating cells) might be responsible for relapse by combining self-renewal capacity with dormancy and treatment resistance (2).AML LIC features have long been considered mainly constant and persistent (2–4); in contrast, recent data suggest unsteady features, i.e. under therapeutic pressure, while data in the absence of treatment remain elusive (5, 6). Putative functional plasticity of AML LIC is of major clinical importance as it might enable novel therapeutic options. We previously reported functional plasticity in acute lymphoblastic leukemia (ALL), where we showed that *in vivo* long-term dormant, treatment-resistant ALL cells were able to convert into highly proliferative, treatment-sensitive cells and vice versa (7). Nevertheless, AML and ALL differ severely regarding stem cell biology, and a defined stem cell hierarchy - characteristic for AML - was never proven in ALL. Based on diverse stem cell characteristics, we considered functional plasticity of LIC conceivable in ALL, but hypothesized its absence in AML.

## Results

To test our hypothesis, we studied cells from eight patients with high risk AML or relapse, of different karyotypes, genotypes and clinical histories (Table S1). As clinic-close model system, primary cells were transplanted into immunocompromised mice, and AML patient-derived xenograft (PDX) models established and serially re-passaged (8). AML PDX models were genetically engineered to express luciferase for bioluminescence *in vivo* imaging and mCherry for cell enrichment by flow cytometry; marker expression remained stable over serial re-transplantation and allowed enrichment of minute numbers of PDX AML cells from murine bone marrow (Figure 1A, see supplemental methods for detailed information). As control, two samples (AML-356 and AML-538) were studied without prior genetic engineering.

**Fig. 1.**
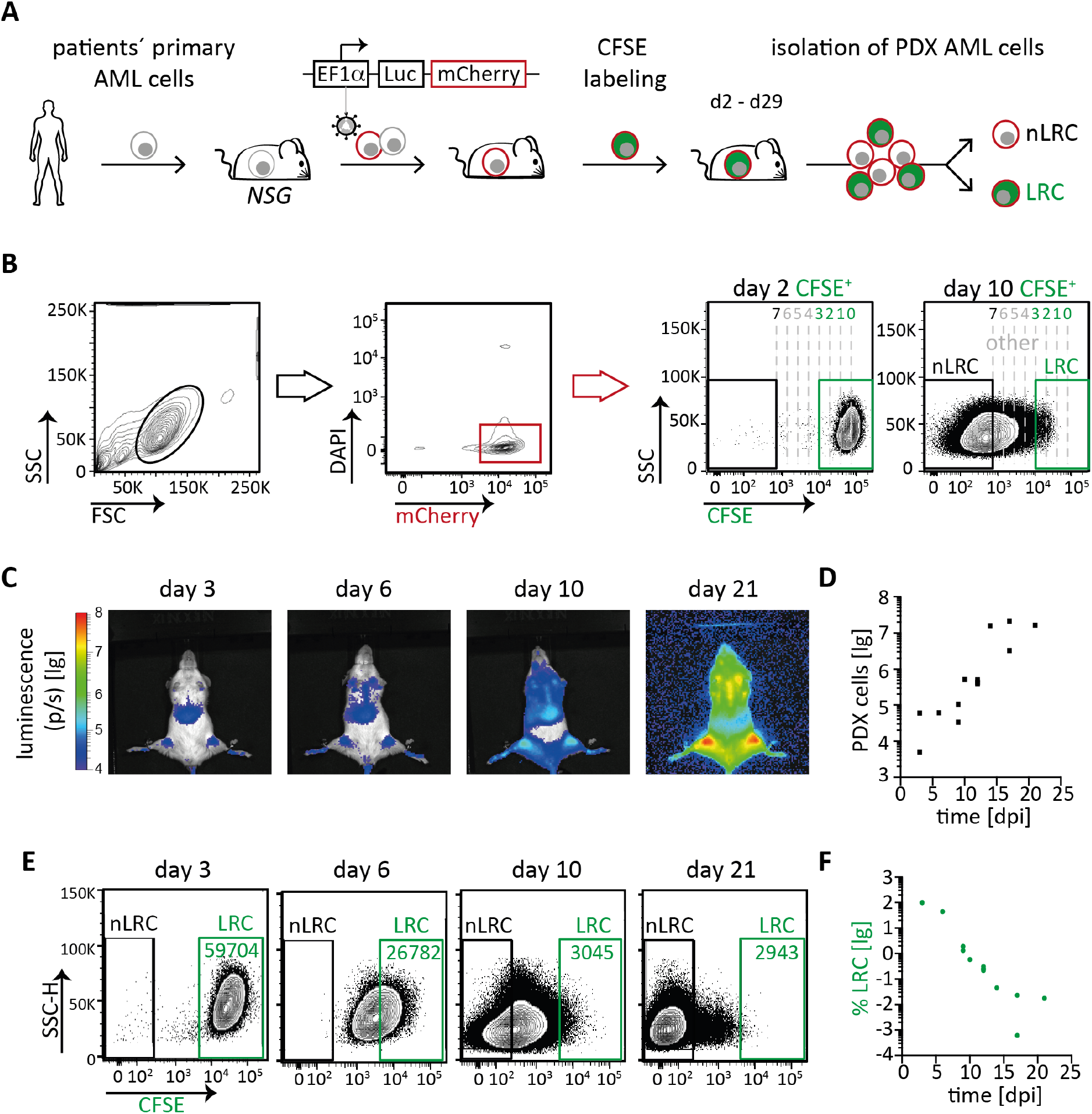
AML PDX cells contain a rare subpopulation of low-cycling cells. **A** Experimental procedure; primary patients’ AML cells were transplanted into NSG mice, resulting PDX cells were genetically engineered by lentiviral transduction, sorted, and amplified. At advanced disease stage, mCherry-positive AML PDX cells were isolated, stained with CFSE, and re-transplanted. At different time points, AML cells were re-isolated from mouse bone marrow, enriched, and CFSE content measured by flow cytometry, to detect CFSE-positive, low-cycling label-retaining cells (LRC), and CFSE-negative, proliferating non-LRC (nLRC). NSG: NOD.Cg-Prkdcscid Il2rgtm1Wjl/SzJ; EF1*α* elongation factor 1-alpha promoter; Luc: enhanced firefly luciferase. **B** Gating strategy; bone marrow cells depleted of murine cells by MACS were gated on (i) leukocytes, (ii) on DAPI-negative mCherry-positive AML PDX cells, and (iii) separated into LRC and non-LRC according to their CFSE content. Maximum CFSE MFI was measured at day two after cell injection or **in vitro** cultivation, and divided by factor two to model cell divisions (dotted lines); upon less than three divisions, cells were considered as low-cycling LRC, upon more than seven divisions as proliferating cells non-LRC; days indicate time after cell injection. **C,D** Growth of AML-393 cells monitored by *in vivo* imaging **(C)** or by quantifying cells re-isolated from mouse bone marrow using flow cytometry **(D)**; each square represents data from one mouse. **E,F** A rare subpopulation of AML PDX cells retains CFSE upon prolonged *in vivo* growth. AML-393 cells from different time points in D were analyzed by flow cytometry for CFSE using the gating strategy described in B; representative dot plots **(E)** and percentage of LRC cells among all isolated PDX cells are shown **(F)**; each dot represents data of one mouse. lg=log10; *See supplemental Figure S1 and S2 for additional data*

AML PDX samples showed heterogeneous growth behavior *in vivo*, with doubling times varying by more than factor three between samples, resulting in variable time to overt disease in mice (Figure S1AB). When PDX cells were re-isolated from murine bone marrow, mCherry expression enabled unbiased enrichment of AML PDX cells, independently from otherwise putatively subpopulation-restricted surface markers on AML cells (Figure 1B) (7, 9). Re-isolation of PDX cells revealed that homing was heterogeneous between samples, as 0.01 to 1% PDX cells could be re-isolated from mice early after transplantation (Figure S1C). The frequency of LIC, as determined in limiting dilution transplantation assays, varied by factor 10 between samples (Figure S1D and Table S2). Thus, our AML PDX cohort of aggressive samples displayed major *in vivo* functional inter-sample heterogeneity, reflecting the known phenotypic heterogeneity of AML (10).

To track *in vivo* proliferation of AML cells from individual samples, PDX cells were stained with CFSE, a dye which is not metabolized in eukaryotic cells, but decreases upon cell divisions and indicates proliferation (11). CFSE records a cell’s proliferative history rather than providing a snapshot on the cell’s proliferative state at a given moment. CFSE content was measured by flow cytometry at different time points following injection into groups of mice. In accordance with an increase in both, leukemic burden and numbers of re-isolated cells (Figure 1CD), most AML PDX cells entirely lost CFSE within days of *in vivo* growth, indicating high proliferative activity in the majority of cells (Figures 1EF). However, a minor subpopulation of cells retained CFSE over several weeks, indicating a low-cycling, putatively dormant phenotype (Figures 1EF and S2AB). These label-retaining cells (LRC) were reliably present in 7/8 samples tested (Figures 1EF and S2AB). As exception, a single sample of a child with deadly AML relapse had entirely lost the LRC population between day 7 to 15 (Figure S2C), again highlighting the known heterogeneity of AML (10). Nevertheless, in the majority of cases, our data reveal heterogeneity of *in vivo* growth behavior within individual AML PDX samples, including a subpopulation of low-cycling LRC.

Given the long-known link between dormancy and chemoresistance (12), we next compared drug response between low-cycling LRC and high-cycling non-LRC. Groups of mice engrafted with CFSE-labeled cells were treated with a chemotherapeutic regimen mimicking “7+3” induction therapy (1), consisting of DaunoXome and Cytarabine (Figure 2A). *in vivo* treatment clearly diminished tumor burden by at least one order of magnitude (Figures 2B and S3A). Interestingly, low-cycling LRC revealed decreased sensitivity towards systemic treatment compared to high-cycling non-LRC in all samples tested; as net effect, the relative proportion of LRC was enriched in treatment surviving cells (Figures 2CD and S3B). Thus, low-cycling LRC show increased resistance against conventional chemotherapy *in vivo* compared to high-cycling non-LRC.

**Fig. 2.**
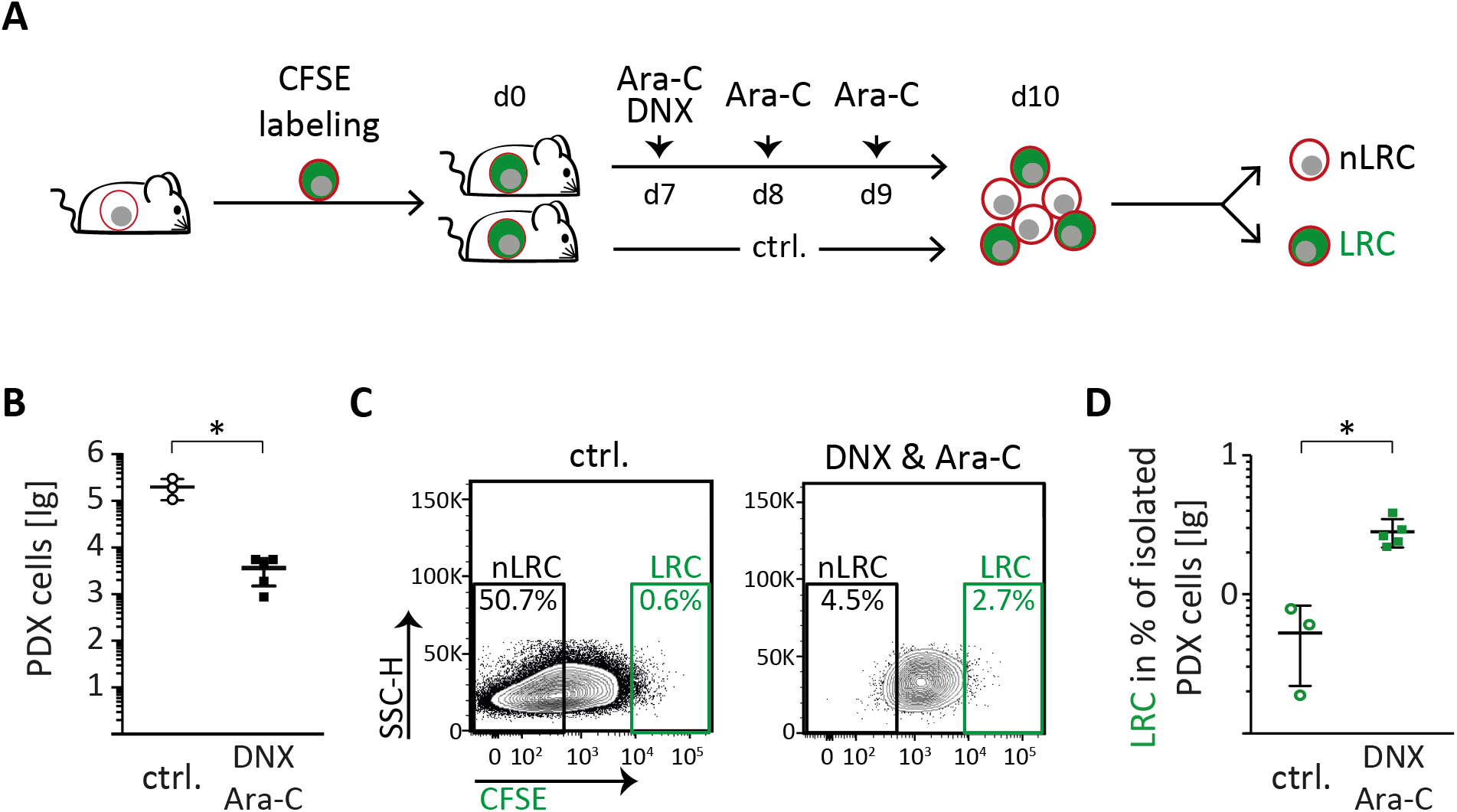
Low-cycling AML PDX cells are chemotherapy resistant *in vivo*. **A** Experimental procedure; groups of mice were injected with CFSE labeled AML-393 cells and treated with PBS (ctrl.; n=3) or a combination of 20 mg/kg DaunoXome® (DNX) on day 7 and 150 mg/kg cytarabine (Ara-C) on days 7 to 9 (n=5); PDX cells were re-isolated from murine bone marrow on day 10 and analyzed as described in Figure 1B. **B** Total number of isolated PDX cells of control and treated mice are shown as mean+/SD; each dot/square represents one mouse. **C,D** Representative dot plots **(C)** and percentage of LRC cells among all isolated PDX cells as mean +/− SD of control and treated mice are shown **(D)**; each dot/square represents one mouse.* p<0.01 by two-tailed unpaired t-test. lg=log10; *See supplemental Figure S3 for additional data*.

We next asked whether the low-cycling state represents a constant, permanent characteristic of a defined AML subpopulation and performed re-transplantation experiments. Low numbers of sorted LRC and non-LRC were re-injected into secondary recipient mice in limiting dilutions (Figures 3A and S4A). Interestingly, both LRC and non-LRC gave rise to leukemia upon re-transplantation, indicating that both subpopulations contained LIC (Figures 3B and S4B). As leukemia development was highly similar in mice transplanted with either LRC or non-LRC, low-cycling LRC must have converted into an actively proliferative state. Furthermore, we found identical LIC frequencies in LRC and non-LRC (Figure 3C, S4C and Table S3), strengthening previous findings (13). These data indicate that LIC reside in both, the low-cycling LRC and high-cycling non-LRC compartment, indicating heterogeneity in proliferation dynamics within the AML LIC pool. As low-cycling cells were able to convert into active proliferation, we asked whether the switch would also be vice versa, so that LRC could be replenished from non-LRC. We re-transplanted high cell numbers, which was technically restricted to non-LRC. Upon secondary transplantation, non-LRC gave rise to a clear LRC fraction (Figures 3DE and S4DE), comparable to the one from bulk cells at first transplantation (Figure 1E and S2A), indicating that high-cycling cells converted to a low-cycling phenotype. These experiments revealed major functional plasticity of AML LIC, and the ability to change their proliferation rate upon changes in external stimuli, such as re-transplantation.

**Fig. 3.**
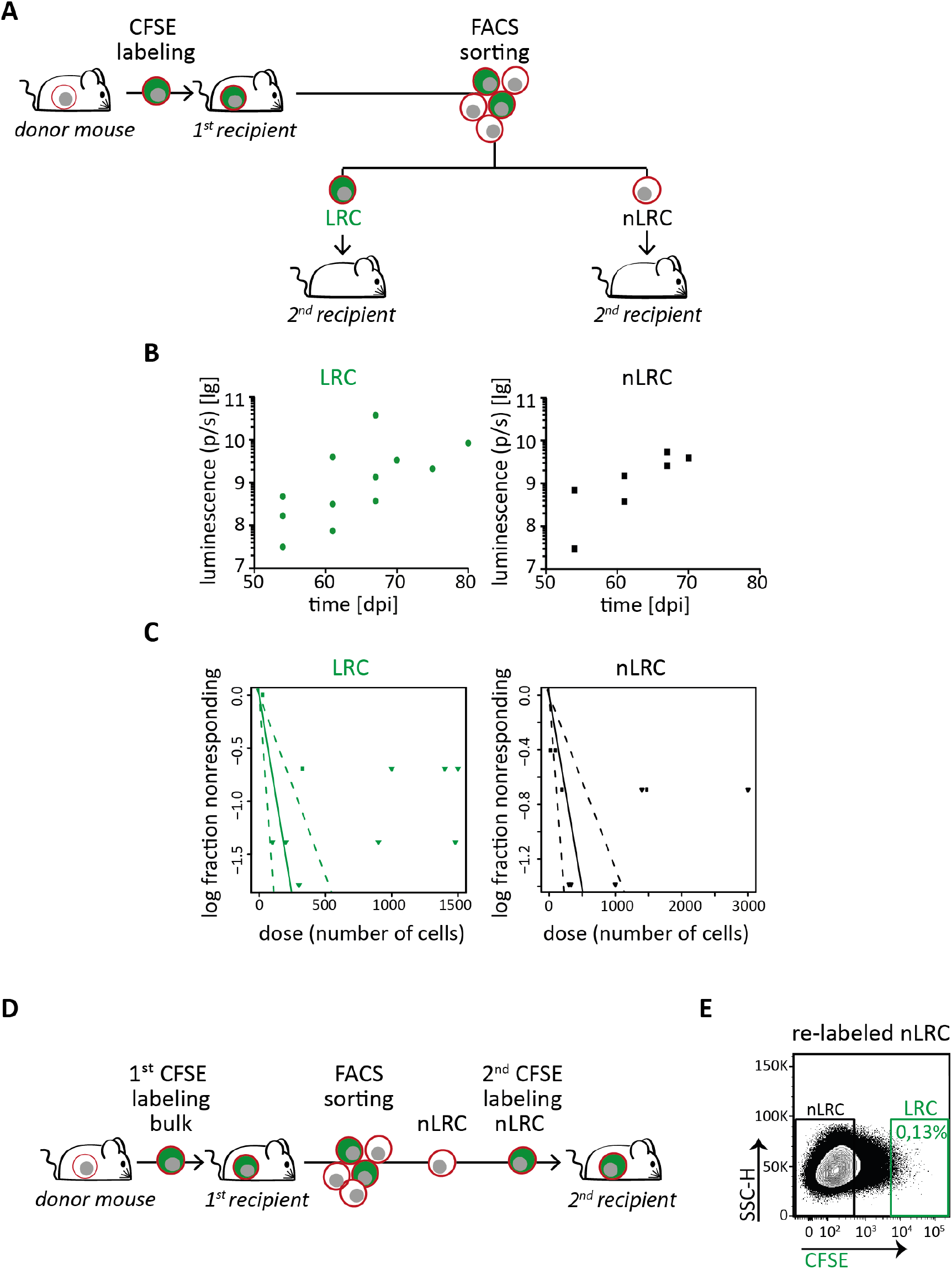
AML PDX cells display reversible growth behavior, independently from stemness potential. **A** Experimental procedure; AML-393 cells were isolated from advanced disease donor mice (n=5 in three independent experiments), labeled with CFSE, and re-transplanted into first recipient mice. Ten days after injection, cells were re-isolated and sorted into LRC and non-LRC (nLRC) using the gates as described in Figure 1B and re-injected into secondary recipient mice. **B** Secondary recipient mice receiving either 300 LRC or 300 non-LRC (n=5) were monitored by *in vivo* imaging. **C** LRC and non-LRC were re-injected into secondary recipient mice (n=38) in limiting dilutions at numbers as indicated in Table S3. Positive engraftment of PDX cells was determined by *in vivo* imaging and/or flow cytometry. LIC frequency was calculated using the ELDA software and is depicted +/− 95% confidence interval. No statistically significant difference between LIC frequency of LRC and non-LRC was found according to chi-square test (p=0.0638). **D,E** From the first recipient mouse harboring CFSE stained cells, 10^7^ non-LRC were isolated at day 21, re-stained with CFSE and injected into secondary recipients; cells were re-isolated 10 days later and LRC were quantified using gates as described in Figure 1B. The experiment is technically unfeasible for LRC as the high number of 10^7^ cells cannot be generated. lg=log10; *See supplemental Figures S4 and Table S3 for additional data*.

## Conclusions

Taken together, our data show that AML contains a rare fraction of low-cycling, treatment resistant LIC which is functionally plastic; any given AML LIC might temporarily adopt a low-cycling LRC phenotype or switch to a rapidly proliferating non-LRC phenotype, depending on, e.g., the surrounding environment. Even the highly aggressive AML samples used in this study harbor the potential to adopt a proliferative phenotype associated with treatment response. Unexpectedly, we detected an identical functional plasticity in AML as in ALL (7), despite that both diseases differ substantially regarding their stem cell biology; ALL never revealed a stem cell hierarchy as proven in AML. In contrast to ALL, AML plasticity comes at a major surprise, as AML LIC were long considered a defined constant subpopulation that persistently and unchangeably combines the adverse characteristics self-renewal, dormancy, and chemo-resistance (2). In our experiments, LIC were not enriched in the LRC fraction, suggesting that dormancy and stemness are not consistently linked in AML. In addition to the known constant, presumably deterministic factors defining stemness, AML stem cells appear to be regulated by additional, transient and putatively stochastic factors (14).

Our data strengthen recent reports demonstrating that chemotherapy can trigger cell cycle entry of AML stem cells (5, 6). In line with these results, our data show that AML LIC respond to external stimuli regarding their proliferative behavior. We here provide the first formal evidence that AML LIC are functionally plastic in the absence of chemotherapy; low proliferating AML LIC convert into rapidly proliferating cells to repopulate the malignancy upon re-transplantation and importantly, highly proliferative LIC can return into a low-cycling state, associated with therapy resistance.

Our data strongly support the concept that retrieving AML LIC from their protective bone marrow niche might sensitize them towards, e.g., conventional chemotherapy (4, 6, 15). The discovered reversibility of the low- and high-cycling phenotypes implicates a direct need to identify factors responsible for AML plasticity, in addition to known microenvironment-derived regulators such as G-CSF (2). As attractive therapeutic concept, inhibition of the reversible low-cycling state might enable overcoming treatment resistance, remove AML LIC, prevent relapse, and ultimately increase patients’ prognosis.

## Supporting information

Supplemental Information_Ebinger et al.

## ACKNOWLEDGEMENTS

We thank Liliana Mura, Fabian Klein, Maike Fritschle, Annette Frank and Miriam Krekel for excellent technical assistance; Markus Brielmeier and team (Research Unit Comparative Medicine) for animal care services; Claudia Baldus and Lorenz Bastian (Divison of Hematology and Oncology, Charité Universitätsmedizin Berlin, Germany) for kindly providing primary cells of AML-538, and Maya C. André and Martin Ebinger (Department of Pediatric Hematology/Oncology, University Children’s Hospital Tuebingen) for kindly providing pediatric AML PDX samples.

