## Supplemental Information_Ebinger et al. for "Functional plasticity and reversible growth behavior of patients’ acute myeloid leukemia stem cells growing in mice"

by Ebinger et al.

### This pdf contains

- **Supplemental Methods**
- **Supplemental Tables (3)**
- **Supplemental Figures (4)**

### Supplemental Methods

**Patients' acute myeloid leukemia (AML) cells.** Bone marrow (BM) or peripheral blood (PB) samples from adult AML patients were obtained from the Department of Internal Medicine III, Ludwig-Maximilians-Universität, Munich, Germany, during the years 2012 and 2014. Specimens were collected for diagnostic purposes before start of treatment. Written informed consent was obtained from all patients. The study was performed in accordance with the ethical standards of the responsible committee on human experimentation (written approval by the Research Ethics Boards of the medical faculty of Ludwig-Maximilians-Universität, Munich, number 068-08 and 222-10) and with the Helsinki Declaration of 1975, as revised in 2000. AML-538 was kindly provided by Claudia Baldus and Lorenz Bastian (Charité Universitätsmedizin Berlin, Germany). Pediatric AML PDX samples were a gift from Maya C. André and Martin Ebinger (University Children's Hospital Tuebingen, Germany), and were described previously (1). Genetic profiling of AML PDX and primary AML samples was performed by Maja Rothenberg-Thurley and Klaus H. Metzeler, as described previously (2).

**The patient derived xenograft (PDX) mouse model of patients' AML.** Xenotransplantation and establishing AML PDX cells in NSG mice (NOD-scid-gamma, The Jackson Laboratory, Bar Harbour, ME, USA) was performed as described previously (3). In the study presented here, only AML PDX cells were applied that re-engrafted in NSG mice over several passages, and lead to a BM chimerism above 90% hCD33+ hCD45+ cells within 16 weeks after transplantation. These requirements precluded the use of primary patient cells, slow engrafters, low engrafters, or samples without the capacity to re-engraft; therefore, the PDX cohort used in this study is enriched for highly aggressive samples. All animal trials were performed in accordance with the current ethical standards of the official committee on animal experimentation (written approval by Regierung von Oberbayern,; 55.2-1-54-2531-95-10, ROB-55.2Vet-2532.Vet02-15-193, ROB-55.2Vet-2532.Vet02-16-7 and ROB-55.2Vet-2532.Vet03-16-56). Accuracy of sample identity was verified by repetitive finger printing using PCR of mitochondrial DNA (4).

**Lentiviral transduction of AML PDX cells and enrichment of transgenic cells.** *In vitro* culture, lentiviral constructs, transduction, and sorting of transgenic PDX cells were performed as described previously (3). In the study presented here, cells were transduced with a construct expressing enhanced firefly luciferase and mCherry in equimolar amounts (pCDH-EF1$\alpha$-eFFly-mCherry, available via Addgene #104833). Initial lentiviral transduction efficiencies were between 1% and 30% (AML-346 30%, AML-372 1%, AML-388, 2% AML-393 12%, AML-491 11%, AML-579 1%). PDX cells were sorted using a FACSAria III (BD Biosciences, Heidelberg, Germany) to reach a purity of more than 95% of mCherry-positive cells. As control, two AML PDX samples without transgenic expression of firefly luciferase and mCherry were applied (AML-356, AML-538).

**Bioluminescence *in vivo* imaging (BLI).** BLI and quantification of tumor burden was performed as described previously (3, 5).

**Labeling of PDX cells with carboxyfluorescein succinimidyl ester (CFSE).** Labeling of PDX cells with CFSE was performed as described previously (5). In brief, AML PDX cells were isolated from mice with advanced disease stage, indicated by a BM chimersim of more than 90% mCherry-positive PDX cells. Cells were labeled with CFSE *ex vivo* (Life Technologies, Carlsbad, CA, USA) according to manufacturer's instructions, washed in PBS, and injected into next recipient mice ($10^7$ CFSE-positive PDX cells per mouse). The procedure resulted in CFSE positivity of well above 98% of PDX cells, as validated by flow cytometry. As AML PDX cells are heterogeneous in size, loss of CFSE appears as continuum in flow cytometry, devoid of the distinct peaks known from normal leukocytes.

**Enriching and quantifying PDX and label-retaining cells (LRC) from mouse BM.** To purify AML PDX cells from mouse BM, bones from hip, femura, tibiae, sternum and spine were crushed with a mortar and pistil, cells were washed once in PBS, and filtered through a cell strainer (EASYSTRAINER 70 µM, Greiner bio-one, Frickenhausen, Germany). Murine cells were depleted using magnetic beads according to manufacturer's instructions (Mouse Cell Depletion Kit, Miltenyi Biotech, Bergisch Gladbach, Germany), with the exception that only 100 µl MicroBeads and two columns were used for one mouse BM suspension. Afterwards, only samples without lentiviral constructs were stained with PE mouse anti-human CD33 IgG1 (WM53, Biolegend).As second step, AML PDX cells were analyzed or sorted by flow cytometry by gating on (i) leukocytes in FSC/SSC, and (ii) DAPI-negative living cells and transgenic mCherry-positive or or CD33-positive AML PDX cells using a BD LSRFortessa or FACSAriaIII, respectively (BD Biosciences, Heidelberg, Germany) as shown in Figure 1B. Sorting of LRC and non-LRC was performed with the precision setting "purity" at the FACSAria. To determine the fraction of low-cycling AML PDX cells, LRC were discriminated from non-LRC using CFSE staining as shown in Figure 1B. To quantify LRC, CFSE mean fluorescence intensity (MFI) of CFSE labelled PDX cells either incubated for two to three days *ex vivo* or isolated from a mouse two to three days after injection was measured, which defined the starting condition ("0 divisions"). Day two or three CFSE MFI was divided by factor two to calculate putative CFSE bisections mimicking cell divisions. Cells with a high CFSE signal below three bisections of the maximum CFSE MFI were defined as LRC. Seven CFSE MFI bisections or more were defined as entire loss of the CFSE signal characterizing non-LRC.

***in vivo* treatment trials.** AML PDX cells were injected into groups of mice ($10^7$ CFSE-positive PDX cells per mouse). Seven days after cell injection, mice were treated with a combination of Cytarabine (150 mg/kg dissolved in PBS, i.p.) on days seven, eight, and nine, and one dose of DaunoXome (20 mg/kg i.v.) on day seven. Body weight was measured daily. Tumor burden was monitored on days seven and ten by BLI. At day ten, mice were sacrificed, BM was collected, and AML PDX cells were isolated and analyzed for CFSE label retention as described above.

**Analysis of plasticity of LRC and non-LRC.** AML PDX cells were isolated from a first donor mouse, labeled with CFSE and transplanted into first recipient mice as described above. Ten (AML-393) or 15 (AML-491) days after injection, AML PDX cells were re-isolated and purely sorted into LRC and non-LRC fractions as described above (see also scheme in Figures 3 and S4). Limiting dilutions of sorted cells were re-injected into secondary recipient mice (between 30 and 3000 cells per mouse, see Table S3). Tumor outgrowth was analyzed by BLI and compared between the groups. Engraftment was determined by positive bioluminescence *in vivo* imaging signal, analysis of hCD33+ cells in peripheral blood (PB), and/or analysis of hCD33+ cells in BM by FACS staining. If no AML PDX cells were detectable within 150 days after injection via BLI, in PB or in BM, mice were counted as non-engrafters. LIC frequencies were determined according to Poisson statistics, using the ELDA software application (http://bioinf.wehi.edu.au/software/elda/)(6).
To determine if highly proliferative cells convert into low-cycling cells, non-LRC from a primary recipient mouse were isolated at day 21 (AML-393) or 16 (AML-491), sorted, re-labeled with CFSE, and re-injected into secondary recipient mice ($10^7$ CFSE+ AML-393 or $10^6$ CFSE+ AML-491 cells per mouse). Ten (AML-393) or 15 (AML-491) days after injection, distribution of LRC and non-LRC was analyzed, and compared to the distribution within first recipient mice. For this analysis, many cells are needed for the re-injection into secondary recipient mice. The minute numbers of LRC that can be re-isolated after ten days from first recipient mice cannot be enriched from secondary recipient mice after re-transplantation; therefore, it is technically unfeasible to perform this analysis with the LRC fraction of cells.

**Statistics.** Statistical analyses were calculated using GraphPad Prism 6 software. Two-tailed unpaired t-test was applied to evaluate differences after drug treatment. ELDA software was used to test differences in LIC frequency by chi-square test.

### Supplemental Tables

**Table S1: Clinical characteristics of AML patients.** .

| Sample | disease stage[1] | age[1] [years] | sex | cytogenetics | mutations[2] |
|---|---|---|---|---|---|
| **AML-346** | R1 | 1 | f | int. del(5q)(13q) | c-Kit |
| **AML-356** | R1 | 5 | m | *ND* | *ND* |
| **AML-372** | R1 | 42 | m | complex, incl. -17 | KRAS, TP53 |
| **AML-388** | ID | 57 | m | KMT2A-AF6 | KRAS, CEBPZ |
| **AML-393** | R1 | 47 | f | KMT2A-AF10 | BCOR, KRAS |
| **AML-491** | R1 | 53 | f | del(7)(q2?1) | DNMT3A, BCOR, NRAS, KRAS, ETV6, PTPN11, RUNX1 |
| **AML-538** | R1 | 68 | f | CN | DNMT3A, IDH1 |
| **AML-579** | R1 | 51 | m | CN | NPM1, FLT3-ITD, DNMT3A, IDH1 |

[1]when the primary AML sample was obtained; [2]mutations detected by targeted re-sequencing in PDX cells; ID = initial diagnosis; R1 = 1st relapse; int = interstitial; del = deletion; CN = cytogenetically normal; f = female; m = male; ND = not determined

**Table S2: LIC frequencies of different AML PDX samples.** (related to Figure S1D)

| Sample | # of cells* | # of mice injected / engrafted | LIC frequency (95% CI) |
|---|---|---|---|
| **AML-372** | 100.000 | 3 / 3 | **1/5.125** (1/1.926-1/13.640) |
| | 30.000 | 3 / 3 | |
| | 10.000 | 3 / 3 | |
| | 3.000 | 3 / 1 | |
| | 1.000 | 3 / 0 | |
| **AML-388** | 72.000 | 1 / 1 | **1/3.665** (1//939-1/14.300) |
| | 24.000 | 1 / 1 | |
| | 21.870 | 1 / 1 | |
| | 7.290 | 1 / 1 | |
| | 2.430 | 2 / 1 | |
| | 710 | 1 / 0 | |
| | 270 | 2 / 0 | |
| | 90 | 1 / 0 | |
| | 30 | 2 / 0 | |
| **AML-346** | 100.000 | 4 / 4 | **1/2.337** (1/898-1/6.093) |
| | 20.000 | 3 / 3 | |
| | 10.000 | 4 / 4 | |
| | 2.000 | 3 / 2 | |
| | 1.000 | 4 / 1 | |
| | 100 | 4 / 0 | |
| **AML-491** | 10.000 | 3 / 3 | **1/1.799** (1/945-1/3.426) |
| | 5.400 | 2 / 2 | |
| | 2.000 | 2 / 1 | |
| | 1.800 | 2 / 0 | |
| | 1.200 | 6 / 6 | |
| | 1.000 | 2 / 1 | |
| | 600 | 5 / 0 | |
| | 200 | 3 / 0 | |
| | 100 | 4 / 0 | |
| **AML-393** | 20.000 | 3 / 3 | **1/507** (1/194-1/1.325) |
| | 2.000 | 3 / 3 | |
| | 666 | 3 / 1 | |
| | 200 | 3 / 2 | |
| | 66 | 3 / 1 | |
| **AML-579** | 72.900 | 1 / 1 | **1/351** (1/77.6-1/1.590) |
| | 24.300 | 2 / 2 | |
| | 7.100 | 1 / 1 | |
| | 2.700 | 2 / 2 | |
| | 900 | 1 / 1 | |
| | 300 | 2 / 1 | |

*Cells from different AML samples were transplanted into recipient mice in limiting dilutions at numbers indicated; bioluminescence *in vivo* imaging, blood measurement or bone marrow FACS staining was performed to determine engraftment; LIC frequency was calculated using the ELDA software; 95% confidence interval (CI).

**Table S3: Identical LIC frequencies of LRC and nLRC.** (related to Figure 3B and S4B)

| Sample | group | # of cells* | # of mice injected / engrafted | LIC frequency (95% CI) |
|---|---|---|---|---|
| AML-393 | non-LRC | 3.000 | 1 / 1 | **1/352** (1/157.1-1/788) |
| | | 1.480 | 2 / 1 | |
| | | 1.400 | 1 / 1 | |
| | | 1.000 | 2 / 2 | |
| | | 330 | 2 / 2 | |
| | | 300 | 2 / 2 | |
| | | 200 | 2 / 1 | |
| | | 100 | 3 / 1 | |
| | | 30 | 3 / 1 | |
| | LRC | 1.500 | 1 / 1 | **1/132** (1/59.2-1/294) |
| | | 1.480 | 2 / 2 | |
| | | 1.400 | 1 / 1 | |
| | | 1.000 | 1 / 1 | |
| | | 900 | 2 / 2 | |
| | | 330 | 2 / 1 | |
| | | 300 | 3 / 2 | |
| | | 200 | 2 / 2 | |
| | | 100 | 2 / 2 | |
| | | 30 | 3 / 0 | |
| AML-491 | non-LRC | 2.000 | 1 / 1 | **1/1080** (1/336-1/3.474) |
| | | 1.200 | 2 / 2 | |
| | | 950 | 1 / 0 | |
| | | 600 | 1 / 0 | |
| | LRC | 2.000 | 1 / 1 | **1/1021** (1/324-1/3.225) |
| | | 1.200 | 2 / 2 | |
| | | 600 | 2 / 0 | |
| | | 200 | 1 / 0 | |

*LRC and non-LRC from first recipient mice were sorted and were transplanted into secondary recipient mice in limiting dilutions at numbers indicated; bioluminescence *in vivo* imaging, blood measurement or bone marrow FACS staining was performed to determine engraftment; LIC frequency was calculated using the ELDA software; 95% confidence interval (CI).

### Supplemental Figures

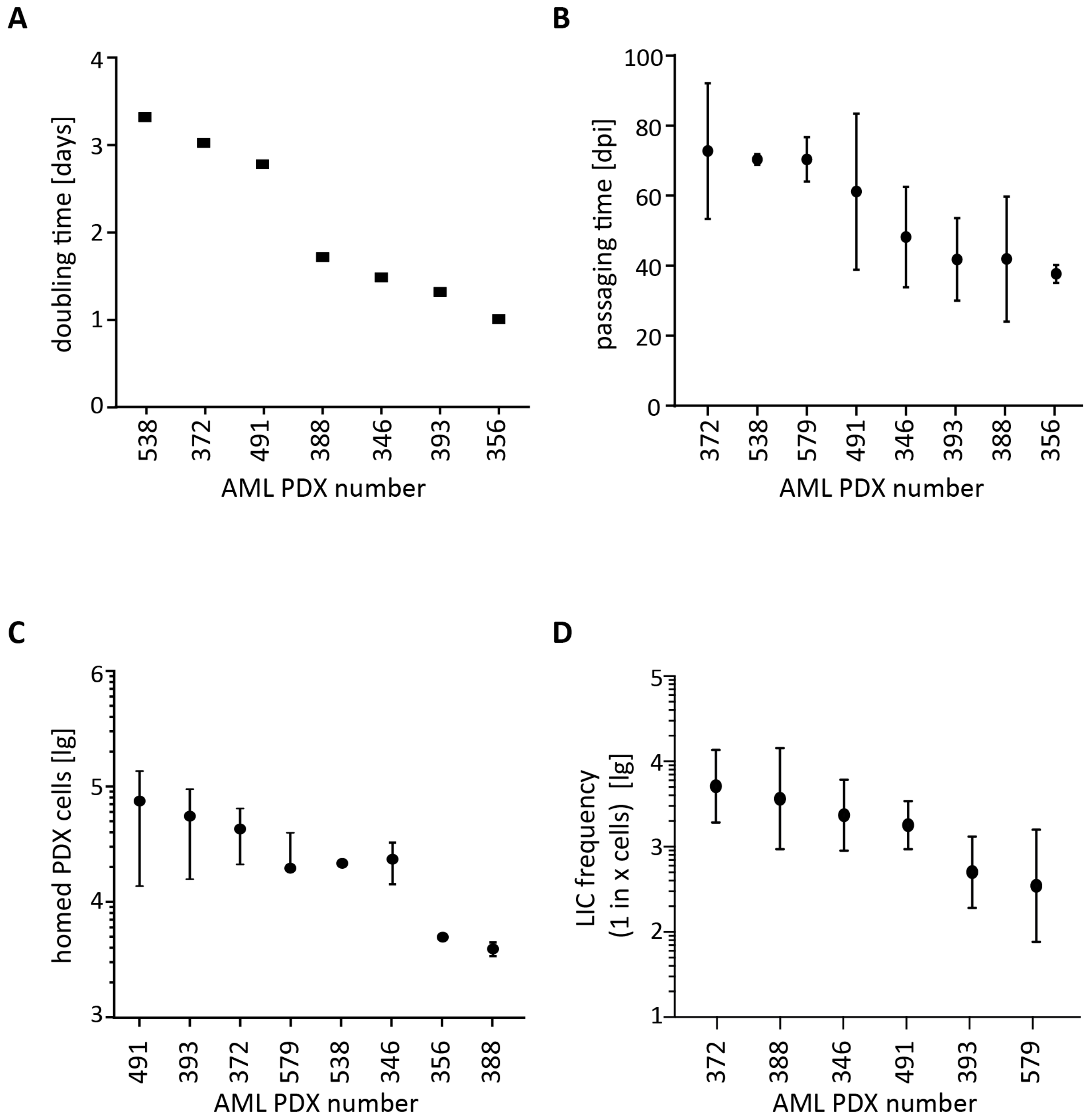


**Figure S 1. AML PDX samples display heterogeneity regarding *in vivo* proliferation, homing and LIC frequency** (related to Figure 1).

**A** *in vivo* doubling times were calculated out of growth curves measured as in Figures 1D and S2.

**B** Passaging times from injection until overt leukemia, $5x10^5$ to $5x10^6$ AML PDX cells per mouse; mean +/- SD of at least 4 and up to 100 mice per sample is depicted. dpi=days post injection.

**C** Number of AML PDX cells homing to the BM was determined 2 or 3 days following injection of $10^7$ cells; mean +/- SD of at least 3 mice is shown, except for AML-538 and AML-356 where a single mouse was analyzed. lg=log10

**D** Bulk cells from different AML samples were transplanted into recipient mice in limiting dilutions at numbers indicated in Table S2. LIC frequency was calculated using the ELDA software and mean +/- 95%CI is depicted. lg=log10

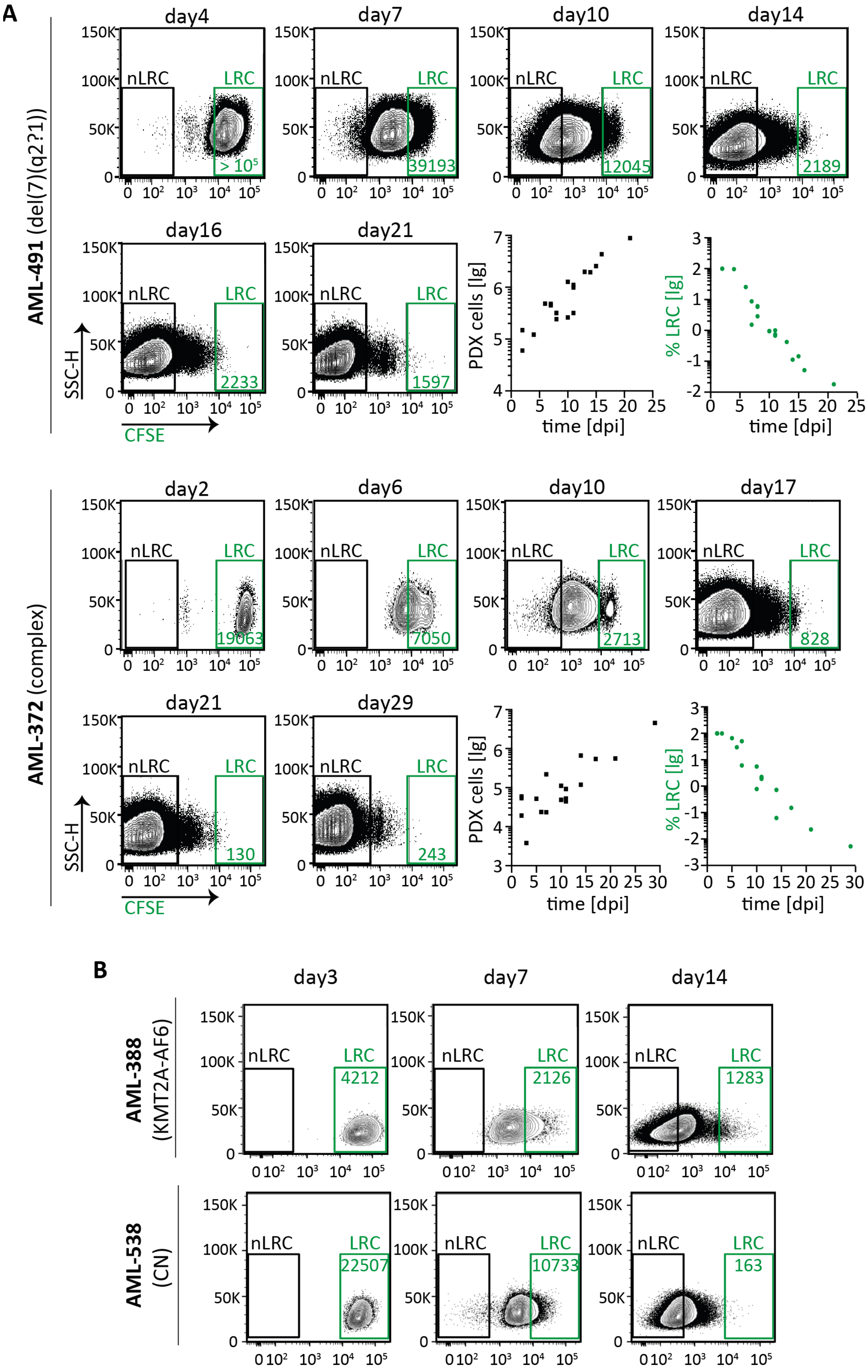
A
AML-491 (del(7)(q2?1))
day4
day7
day10
day14
day16
day21
nLRC
LRC
> 10^5
39193
12045
2189
2233
1597
SSC-H
CFSE
PDX cells [lg]
% LRC [lg]
time [dpi]
AML-372 (complex)
day2
day6
day10
day17
day21
day29
19063
7050
2713
828
130
243
B
AML-388 (KMT2A-AF6)
AML-538 (CN)
day3
day7
day14
4212
2126
1283
22507
10733
163

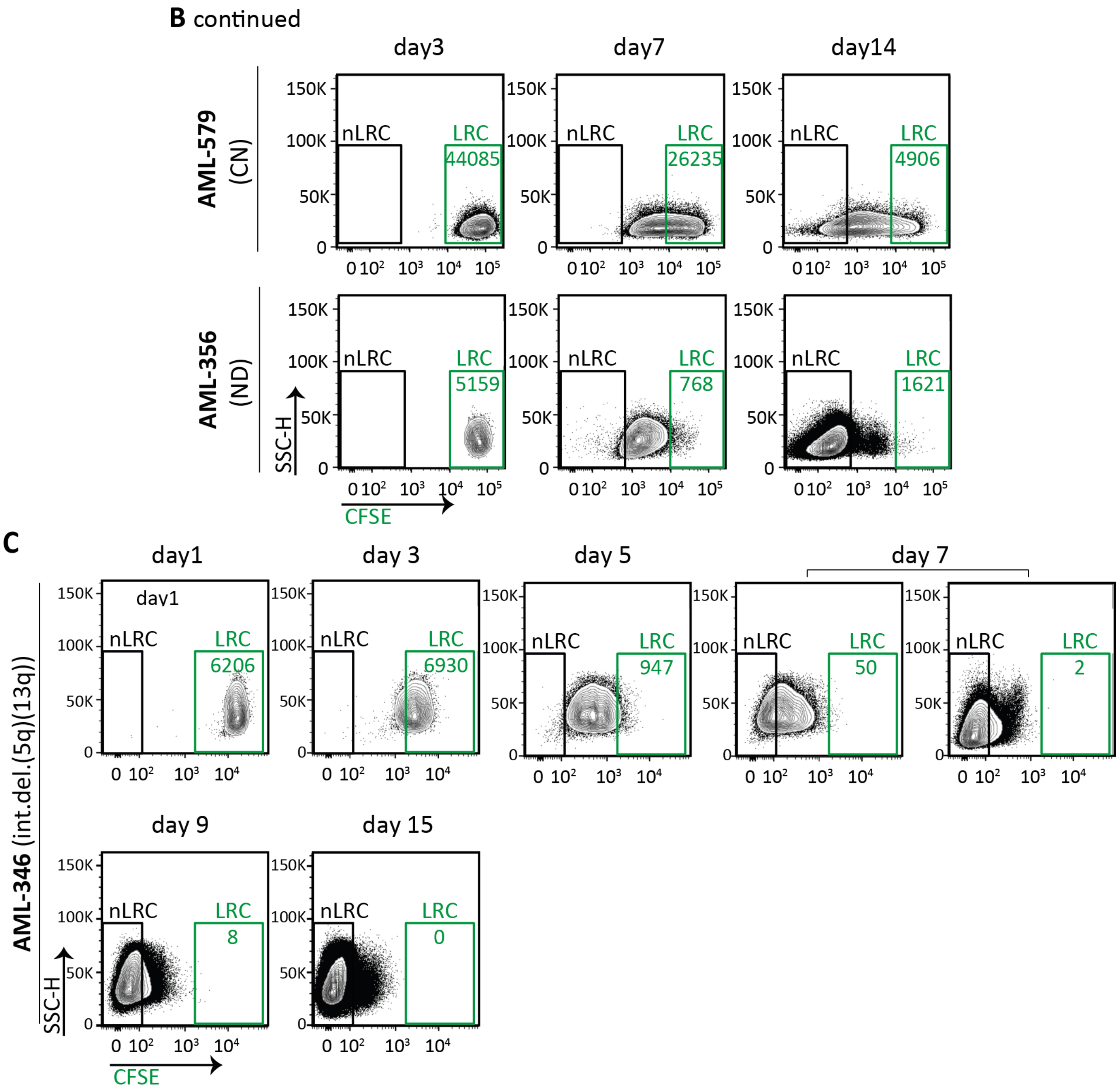


**Figure S 2. AML PDX cells contain a rare subpopulation of low-cycling cells** (related to Figure 1, additional samples).
Experiments were performed and depicted identically as in Figure 1. In brief, $10^7$ CFSE- labeled AML PDX cells were injected into groups of mice; cells were isolated at different time points and analyzed by flow cytometry for CFSE content. FACS plots for representative mice are shown; total number of isolated PDX cells and percentage of LRC cells among all isolated PDX cells are shown in **A**. Total number of mice studied was (**A**) 19 for AML-491, 19 for AML-372, (**B**) 5 for AML-388, 3 for AML-538, 8 for AML-579, 3 for AML-356 and (**C**) 10 for pediatric AML-346. lg=log10

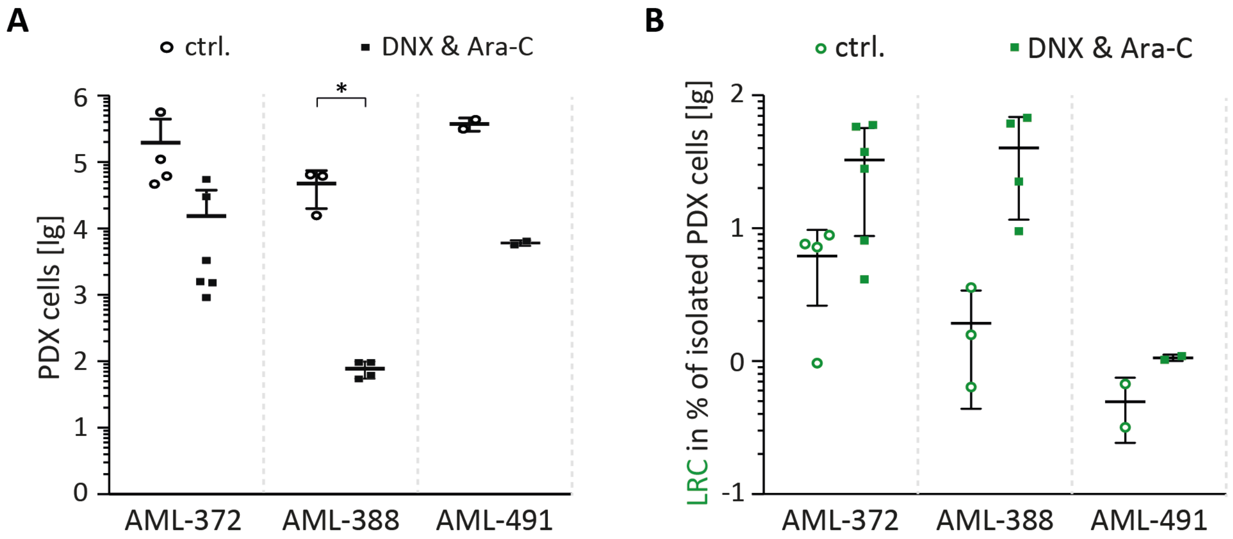


**Figure S 3. Low-cycling LRC display treatment resistance *in vivo*** (related to Figure 2, additional samples).

Experiments were performed and depicted identically as in Figure 2. In brief, groups of mice were injected with CFSE-labeled AML PDX cells and treated with PBS (ctrl.) or a combination of 20 mg/kg DaunoXome® (DNX) on day 7 and 150 mg/kg cytarabine (Ara-C) on days 7 to 9; PDX cells were re-isolated from murine bone marrow on day 10, murine cells depleted by MACS and cells analyzed with flow cytometry (gating see Figure 1B).

**A** shows effect of treatment on bulk PDX cell numbers;

**B** shows effect of treatment on LRC.

Mean +/- SD for AML-372 (n=10), AML-388 (n=7) and AML-491 (n=4) is shown. Each dot/square represents one mouse. * p<0.05 by two-tailed unpaired t-test.

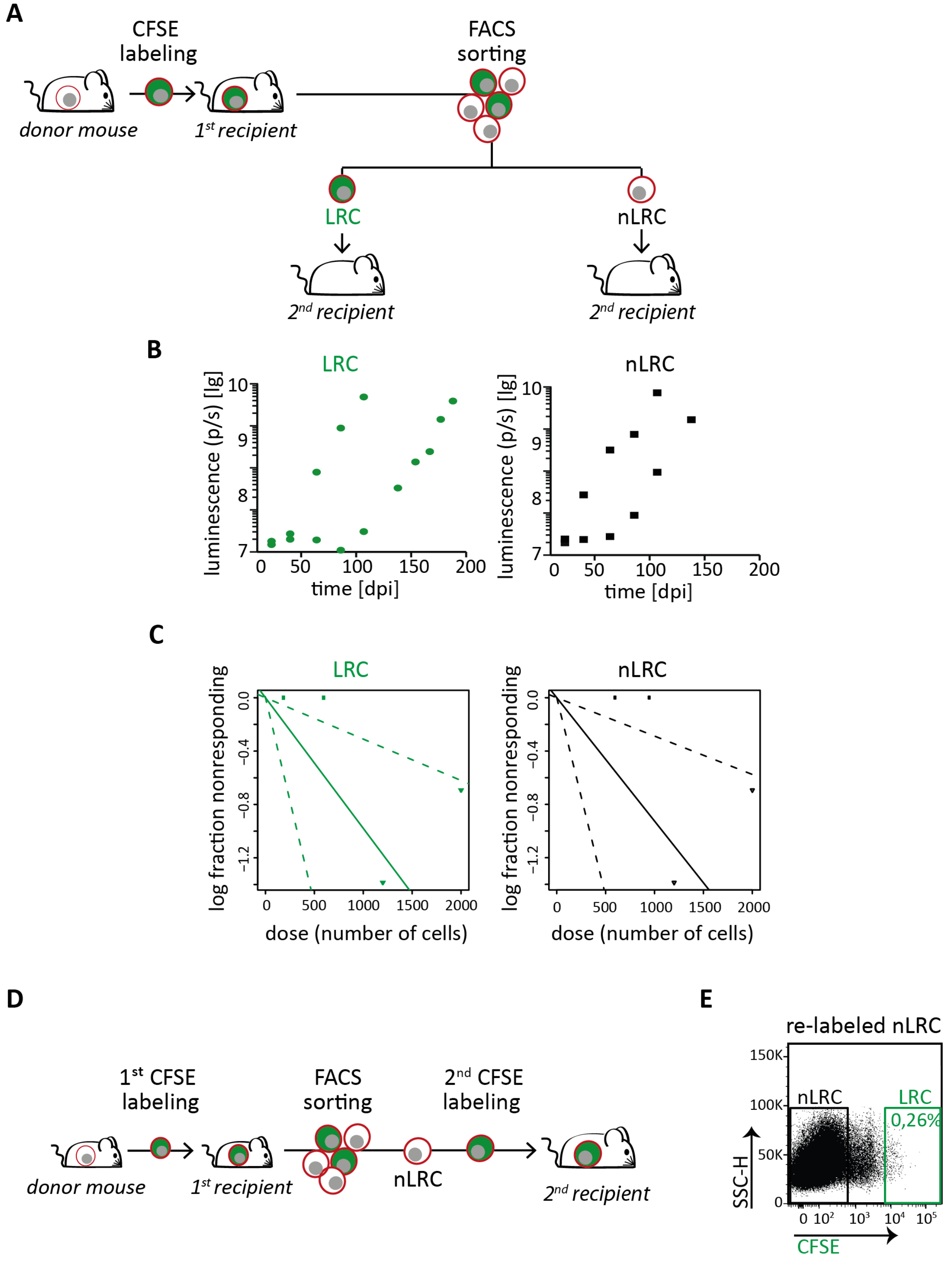


**Figure S 4. AML PDX cells display reversible growth behavior, independently from stemness potential** (related to Figure 3, additional sample AML-491)
Experiments were performed and data depicted identically as in Figure 3;

**A** Experimental procedure; AML-491 cells were isolated from advanced disease donor mice (n=2 in two independent experiments), labeled with CFSE, and re-transplanted into first recipient mice. Sixteen days after injection, cells were re-isolated and sorted into LRC and non-LRC (nLRC) using the gates as described in Figure 1B and re-injected into secondary recipient mice.

**B** Secondary recipient mice receiving either 1200 LRC or 1200 non-LRC (n=4) were monitored by *in vivo* imaging (photons/second).

**C** LRC and non-LRC were re-injected into secondary recipient mice (n=11) in limiting dilutions at numbers as indicated in Table S3. Positive engraftment of PDX cells was determined by *in vivo* imaging and/or flow cytometry. LIC frequency was calculated using the ELDA software and is depicted +/- 95% confidence interval. No statistically significant difference between LIC frequency of LRC and non-LRC was found according to chi-square test (p=0.95).

**D,E** From the first recipient mouse harboring CFSE stained cells, $10^6$ non-LRC were isolated at day 15, re-stained with CFSE and injected into secondary recipients; cells were re-isolated 15 days later and LRC were quantified using gates as described in Figure 1B. The experiment is technically unfeasible for LRC as the high number of $10^6$ cells cannot be generated.

lg=log10; *See supplemental Table S3 for additional data.*
